# Evolutionary implementation of Bayesian computations

**DOI:** 10.1101/685842

**Authors:** Dániel Czégel, Hamza Giaffar, István Zachar, Joshua B. Tenenbaum, Eörs Szathmáry

## Abstract

The Bayesian framework offers a flexible language for the consistent modular assembly of statistical models used by both minds and machines. Another algorithmic domain capable of adaptation in potentially high-dimensional and uncertain environments is Darwinian evolution. The equivalence of their fundamental dynamical equations, replicator dynamics and Bayesian update, hints at a deeper algorithmic analogy. Here we show, based on a unified mathematical discussion of evolutionary dynamics and statistical learning in terms of Bayesian graphical models, that this is indeed the case. Building blocks of Bayesian computations, such as inference in hierarchical models, filtering in hidden Markov models, gradient likelihood optimization, and expectation-maximization dynamics of mixture models, map naturally to fundamental concepts of evolution: multilevel selection, quasispecies dynamics, phenotypic adaptation and ecological competition, respectively. We believe that these correspondences point towards a more comprehensive understanding of flavors of adaptive computation observed in Nature, as well as suggesting new ways to combine insights from the two domains in engineering applications.

## 1 Introduction

The Bayesian framework of performing consistent computations about probabilistic beliefs has become a fundamental language to describe and understand adaptation in the presence of uncertainty [1, 2, 3]. Two factors have recently driven the theoretical development of this machinery: i) the machine learning revolution building on the boom of available data and computational power [4, 5] and ii) the possibility to perform largescale (human and non-human) cognitive experiments that necessitates a modeling framework accounting for trial-to-trial and inter-individual variability in a probabilistic manner [6]. A Bayesian approach, building on the concept of belief update driven by external evidence, seems to provide a modeling framework that possesses the appropriate amount of richness to build machines that represent the environment efficiently by either human designers or by the “blind” processes of inheritance and natural selection [7, 8]. One major advantage of Bayesian models is that they naturally translate between a generative direction that predicts data given current model and the reverse process of inference that updates the model given new data. This allows for online learning, naturally accounting for the optimal level of uncertainty on-the-fly given the amount of data that is available [9].

The fact that many efficient adaptive mechanisms that leverage stochastic information about the external world seemingly converge on the same basic set of Bayesian computations points at the importance of understanding the possible ways of implementing such computations. Within neuroscience, advances have been made in formulating possible implementations of Bayesian computations that are consistent with the constraints given by neural substrates, including locality of both information processing and learning and also the spiking nature of neural computations [10].

In this paper, we explore how different flavors of Bayesian computations can be implemented on a fundamentally different substrate: the substrate of Darwinian replicators. The common overarching theme of such implementations is that abundances of types (ribozymes, genes, cells, organisms, etc. in a biological setting) represent probabilities of hypotheses. In analogy to the updating of probabilities associated to hypotheses based on their ability to predict current data, the abundances of types are updated based on a similar kind of environmental feedback: their fitness in the current environment. This analogy can be formalized by the equivalence of the discrete-time replicator equation and Bayesian update [11, 12]. Bayesian update quantifies how the prior probability *P* (*h*_*i*_) of hypothesis *h*_*i*_ is updated to its posterior probability *P* (*h*_*i*_|*x*), given data *x* and the probability *P* (*x*|*h*_*i*_) of *x* being generated (or predicted) by *h*_*i*_ called the likelihood of *h*_*i*_, as

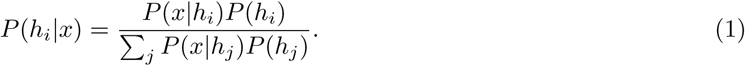

The discrete-time replicator equation has the equivalent algebraic form: the relative abundance of type *i*, denoted by *p*_*i*_ is updated from time *t* to *t* + 1 according to its fitness *f*_*i*_(*x*) in the current environment *x* as

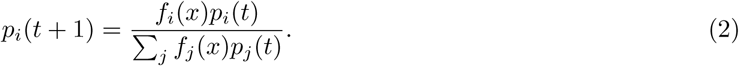

Identifying relative abundances with probabilities and fitness with likelihood shows that these two forms of update are indeed equivalent.

The goal of this paper is to point out how any evolutionary dynamics, natural or engineered, biological or non-biological, fits into a general Bayesian computational framework. Two potential benefits of this approach are i) it might help understanding (macro)evolutionary processes in high-dimensional stochastic environments in normative/computational terms and ii) it can hint at possible ways to combine evolutionary and Bayesian computations, leveraging the advantages of both in a common computational framework [13, 14].

Thanks to recent efforts, many fundamental Darwinian phenomena can now be translated to the language of Bayesian computations, including selection [11, 12], mutation [15] and multilevel evolutionary processes [16]. Here we complement this list with the Bayesian interpretation of phenotypic adaptation and of evolutionary-ecological dynamics.

More importantly, we discuss elementary building blocks of evolutionary and probabilistic models in a coherent narrative, provided by the language of graphical models [3], which, similarly to Feynman diagrams in particle physics, provide an easy-to-manipulate visual representation of relevant aspects of computations, while, just as importantly, hides less relevant ones.

We also believe that one of the most important contributions any theoretical approach can provide is the clear articulation of assumptions behind any argumentation. We therefore attempt to provide exact mathematical formulations wherever possible.

### Building blocks of evolutionary dynamics and statistical learning

In the following, we build up a series of correspondences between simple models of evolutionary dynamics, including that of selection, adaptation, mutation, evolutionary-ecological dynamics and multilevel selection, and Bayesian generative models in a step-by-step manner, highlighting the assumptions we make at each step. Importantly, we also point out what aspects of modeling are not constrained by this theoretical framework.

First and foremost, one of the most unconstrained aspects of this correspondence is the interpretation of environment *x*(*t*) at time *t*. The environment can comprise any number of data points at each time step, and can be governed by any (unknown) generative process. Among others, two possible interpretations are the following: *x*(*t*) might correspond to one single data point summarizing the ecologically relevant dimensions of the actual environment without further specification, or, *x*(*t*) might correspond to a set of points, whose local density stands for the probability/weight of the environment being at that given state. This includes the interpretation of the density of points corresponding to the density of finite resources, one that is assumed when ecological competition is modeled.

Another cornerstone connection between evolutionary and Bayesian models is the interpretation of likelihood as fitness. The likelihood function *P* (*x*|*h*_*i*_), telling us the probability of *x* being generated by hypothesis *h*_*i*_, accounts naturally for an important constraint: adaptation always unfolds under some trade-offs that prohibit unlimited mastery of all possible environments; instead, both generative models and replicator types must give up adeptness in one environment to be more adapted to others. This is formalized in simple mathematical terms as the likelihood function being normalized over possible environments, ∑_*x*_ *P* (*x*|*h*_*i*_) = 1, corresponding to ∑_*x*_ *f*_*i*_(*x*) = 1 for any hypothesis/type.

Given the statistical structure of the environment and the constraint on possible fitness functions described above, replicator types compete to fit the environment in a manner analogous to the competition between hypotheses to predict the statistical structure of the data. Depending on the possible modes and timescales of adaptive mechanisms, minimal dynamical models of evolutionary processes map to different Bayesian models and inference procedures. In particular, two relevant timescales of evolution are i) microevolutionary timescale, where the change in abundance distribution over replicator types is explicitly modeled, ii) macroevolutionary timescale, where only the evolution of phenotypic population average is modeled. These are two complementary building block models of evolutionary dynamics, upon which additional ingredients, such as explicit mutation, ecological competition or multilevel selection, can be added. For simplicity, we refer to the former as selection and the latter as adaptation.

### Quasispecies dynamics and filtering in Hidden Markov Models

Selection by itself does not account for how variation is generated; instead, it assumes that replicator types are fixed and only their relative abundances *p*_*i*_(*t*) change over time. Indeed, as described above, the abundance of those replicator types that provide a better fit of the current environment *x* increases at a higher rate according to the discrete-time replicator equation; equivalently, the posterior probability of those hypotheses that predict the current data better increase at a higher rate, described by Bayesian update. The corresponding graphical model is depicted in Figure 1a. This dynamics is the simplest building block that models selection on a microevolutionary timescale.

**Figure 1:**
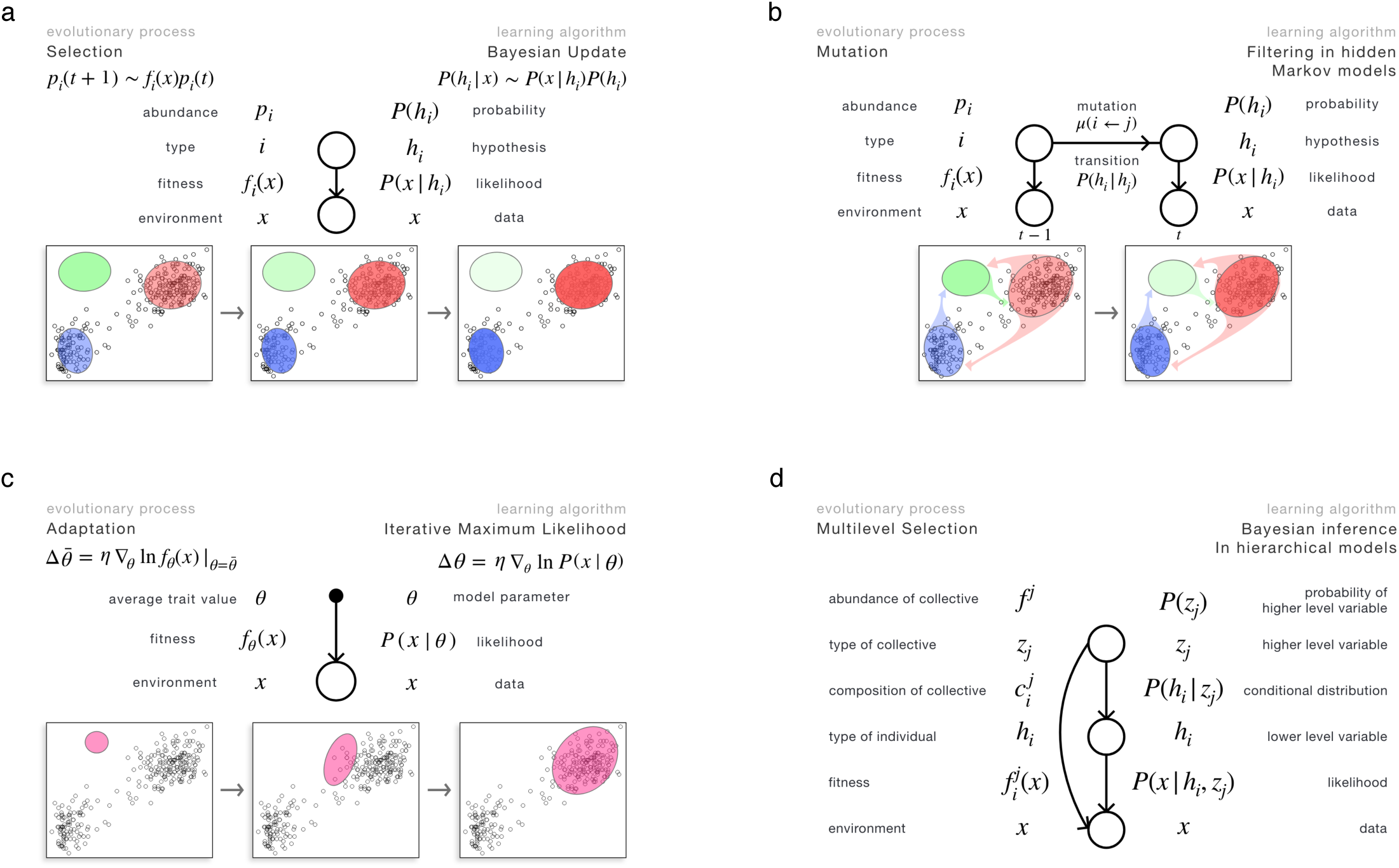
Evolutionary interpretation of elements of statistical learning. a) Replicator equation and Bayesian update. b) Quasispecies dynamics and filtering in hidden Markov models. c) Phenotypic adaptation and gradient likelihood maximization. d) Multilevel selection and inference in hierarchical models. Top: Graphical representation of equivalent dynamical equations and translation table between quantities of evolution and statistical learning. See text for mathematical details. Bottom: Visualization of adaptation dynamics. Environment/data is represented by datapoints; position and orientation of Gaussian contours depict fitness function of types over possible environments/likelihood functions corresponding to different Bayesian generative models. Opacity shows relative frequency of types/posterior probability of generative models. Based on these correspondences, we suggest that the language of Bayesian graphical models might also provide a helpful visualization tool for summarizing structural properties of evolutionary models.

Variation, generated by mutations, can be introduced into the selection-only model on two different timescales. The first approach extends the replicator equation by explicitly characterizing the number of mutation events, modeled by mutation rates *µ*_*i←j*_ from type *j* to type *i*. The evolutionary dynamics is therefore determined by the overall effect of two forces acting on the same timescale: selection and mutation. Mathematically, this simultaneous dynamics can be accounted for by the replicator-mutator equation (cf. eq. (2)),

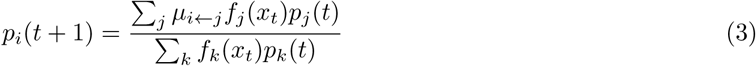

where *µ*_*i←j*_ determines the mutation probability (in unit time) from type *j* to type *i* and therefore is normalized as ∑_*i*_ *µ*_*i←j*_ = 1, and the factor ∑_*k*_ *f*_*k*_(*x*_*t*_)*p*_*k*_(*t*) is the average fitness of the population, responsible for keeping the distribution *p*_*i*_ normalized at all times. Eq. (3) describes how fitness *f*_*i*_(*x*_*t*_) in the current environment, together with the mutation probabilities *µ*_*i←j*_, update the abundance distribution *p*_*i*_. Similarly to the simple replicator equation, information about the environment is transferred to the abundance distribution over time; here, however, this information is filtered through the mutation probabilities, including the diagonal elements *µ*_*i←i*_ that specify the fidelity of replication. As observed by [15], this model of replication and mutation is equivalent to updating the probability distribution over latent hypotheses *h*_*i*_ in hidden Markov models (HMMs). HMMs extend Bayesian update by introducing probabilistic transitions between hidden states (i.e. hypotheses) *h*_*i*_, and they infer the joint distribution *{P* (*h*_*i*_, *t* = 1), *P* (*h*_*i*_, *t* = 2), *…, P* (*h*_*i*_, *t* = *T*)*}* over the hidden states over time given data history *{x*_1_, *x*_2_ *… x*_*T*_ *}*. In order to achieve this, HMMs need two quantities to be pre-determined: hypothesis *h*_*i*_’s likelihood of data, *P* (*x*|*h*_*i*_), just like in the case of simple Bayesian update, and the transition probabilities from *h*_*j*_ to *h*_*i*_, denoted by *P* (*h*_*i*_|*h*_*j*_). This joint probability distribution over hypotheses at all times, *{P* (*h*_*i*_, *t* = 1), *P* (*h*_*i*_, *t* = 2), *…, P* (*h*_*i*_, *t* = *T*)*}*, can be reduced, however, to distributions of interest; if this distribution of interest is the one corresponding to the last timestep, *P* (*h*_*i*_, *t*), dynamically inferring this last distribution given data history is called the *filtering* problem in HMMs, and it is obtained through the dynamics

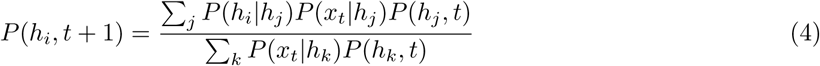

that is indeed equivalent to eq. (3), with the identifications depicted in Figure 1b. Importantly, HMMs can also be cast as Bayesian graphical models, as illustrated in Figure 1b. Note that this dynamics computes the predictive distribution at time *t*+1, corresponding to taking into account data until time *t* and transitions until time *t* + 1; if data in the last timestep *t* + 1 is also integrated, eq. (4) needs a slight modification, corresponding to a less frequently used evolutionary counterpart. One important consequence of this analogy is that an evolutionary search in the vast space of Bayesian model *structures* might be possible within a Bayesian realm. What makes such a procedure practically non-trivial is the need for an efficient (implicit) definition of the mutation kernel between the infinitely many possible model structures. Augmenting this Darwinian-Bayesian dynamics with a parallel search for rare subsolutions provided by recombination is, we believe, another direction worth exploring.

### Phenotypic adaptation and gradient likelihood optimization

The other approach of introducing the effect of mutations, that we refer to as adaptation, corresponds to modeling evolutionary dynamics at a qualitatively longer timescale [17, 18, 19]. It does not explicitly account for mutations, instead, it assumes that a cloud of mutants in the (pheno)type space is always available upon which selection can act, irrespectively of what the generative mechanism of this variation is. It also shifts the attention from tracking the abundance distribution of all types over time to simply following the population average of some relevant phenotypic traits. As an effect, in simple adaptive scenarios (with the explicit assumptions delineated below), the population average of relevant traits, denoted by 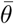, evolves according to a gradient ascent on the (log) fitness landscape:

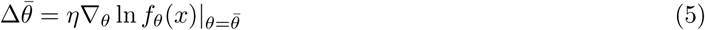

where 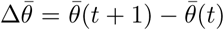 is the change in the population average trait value (that can be a vector of any dimensionality), *f*_*θ*_(*x*) is the fitness of individuals with trait value *θ* given environmental state *x*, and *η* is the rate of adaptation. Indeed, the Price equation [17], in its simplest form (assuming no intergenerational transmission bias), relates the change in population average 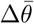 and the covariance between fitness and trait value cov(*θ, f*) as 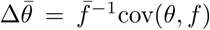, where 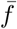 is the average fitness in the population. Assuming a deterministic fitness function *f*_*θ*_ once the environment *x* is fixed, an isotropic cloud of mutants in the phenotype space, and that the variance in trait values is small enough to approximate the fitness linearly around the population average by 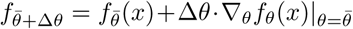, the Price equation implies (see Materials and Methods) the dynamics given by eq. (5) with rate of adaptation being equal to the variance of trait values projected to the direction of fitness gradient, *η* = var(*θ*_||_).

This dynamics, assuming the above mentioned normalization constraint on fitness ∑_*x*_ *f*_*θ*_(*x*) = 1, is equivalent to the gradient optimization of the (log) likelihood function,

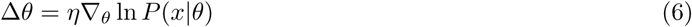

with learning rate *η*. The corresponding graphical model is shown in figure 1c. Crucially, the connection between the two descriptions is given by the identification of fitness and likelihood again. In this model of macroevolutionary adaptation, new types can and do arise; however, it is only the phenotypic population average that this description accounts for. Furthermore, the rate of adaptation *η* is given by the phenotypic population variance that i) might change over time, and ii) is not modeled here, in accordance with the dynamical insufficiency of the Price equation [20].

### Multilevel selection and Bayesian inference in hierarchical models

When the replication of different replicator types are partially but not fully synchronized and groups of replicators inherit information regarding the identity of the group, an effective description of the system is provided by multilevel selection theory (MLS) [21]. Biological examples, where selection is possibly nonnegligible at multiple levels, include genes within protocells, reproducing organelles in the eukaryotic cell, or individuals in a social insect colony. MLS decomposes the full effect of selection hierarchically to selection acting between and within groups. Partially synchronized replicators form a necessary intermediate step towards new emerging units of evolution, i.e., transitions in individuality, that is, in turn, a main mechanism responsible for increasing complexity in evolutionary systems. Understanding MLS is therefore of crucial importance regarding any, natural or engineered, open-ended Darwinian system.

MLS acts on a hierarchical population of replicators: types of individuals *h*_*i*_ are grouped into types of collectives *z*_*j*_. A complete static description of the population is given by the abundance and composition of collectives (for further details, see [16]). The composition of collective *z*_*j*_, denoted by 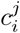, normalized as 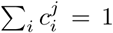, quantifies the relative abundance of individual level replicators within collectives of type *z*_*j*_. The total abundance of collective *z*_*j*_, measured as the total abundance of individual-level replicators within collectives of type *z*_*j*_, is denoted by *p*^*j*^ and normalized as ∑_*j*_ *p*^*j*^ = 1. These two sets of relative abundances, the compositions of collectives 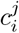 and their abundance, *p*^*j*^, describe the the composition of the hierarchical (two-level) population completely. From these two quantities, a third one, the total abundance of individual-level replicators of type *h*_*i*_ being part of collective *z*_*j*_, denoted by 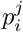, can be calculated as 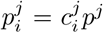. This quantity connects the two population levels in a sense that the abundance distribution of any level can be computed by summing over the other level: at the level of collectives, the abundance distribution, as we have seen, is given by 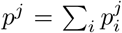; at the level of individuals, the abundance of type *h*_*i*_, *p*_*i*_, is obtained by adding up the abundances of type *h*_*i*_ being in any collective, 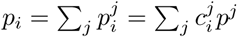.

These quantities above describe the composition of the population at one time instance. How does selection change this hierarchical population over time? According to the discrete-time replicator dynamics, abundances change proportionally to their fitness. Here, however, the fitness of an individual-level type *h*_*i*_ might very well depend on the collective *z*_*j*_ it is part of; we denote this fitness by 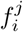. We also allow this fitness to depend on the environment *x*. The replicator dynamics, tracking the abundance of individuals of type *h*_*i*_ being part of collectives *z*_*j*_, then reads as

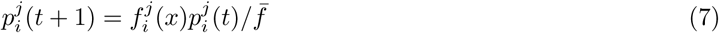

where the average fitness, 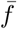, is calculated as 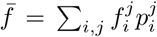. Tracking the abundance distribution at any level is then possible by summing over the other level, as discussed above. This, however, does not mean that the dynamics at the two levels are decoupled, for example, at the level of individuals, 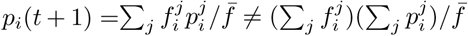.

Crucially, fitnesses and abundances are assigned to types that connect the two levels, namely, to individuals of type *h*_*i*_ that are part of collective *z*_*j*_. The replicator equation acts on these quantities, evolving the multilevel population in time. This conceptualization of MLS allows for relating multilevel evolutionary dynamics to hierarchical Bayesian computations over multivariate distributions: these two hierarchical dynamics are structurally equivalent, with the identified quantities listed in Figure 1d. Note that this analogy can be extended to arbitrary number of levels, see [16] for details.

An important aspect of multivariate Bayesian models is that it is possible to identify a parametrization-independent backbone of the model in terms of conditional independence relations between variables. This allows for distinguishing between model structure (topology) and parameters, a necessary step towards modeling causality [22]. As discussed in [16], conditional independence relations imposed on the evolutionary implementation of the corresponding Bayesian model have a well-defined intuitive meaning, too: it corresponds to freezing compositions at various levels of the hierarchical population.

Another fundamental feature of hierarchical models is their complexity. Finding the probabilistic model with optimal complexity given data is a non-trivial task that can be approached from many directions. The Bayesian approach is to gauge the model’s performance averaged over all possible parameter settings of hidden variables [23]. When comparing two models, possibly having different number of variables and different number of hierarchical levels, one has to favor the one that predict data better on average. Interestingly, this Bayesian agenda translates to a general and simple evolutionary interpretation: if the average fitness of a collection of replicators is higher when they are grouped together in a collective compared to the case when they are “free”, a new level emerges. Although this model of evolutionary transition of individuality [24, 25] is rather simplistic in terms of dynamics, it is not in terms of structure, and we believe that this similarity is quite remarkable, having the potential to be refined later on to understand and design complexifying evolutionary systems.

### Ecological competition and expectation-maximization in mixture models

Selection and adaptation, as well as the combination of the two, that is necessary to describe evolutionary-ecological interaction of species, form the basis of relating more complex evolutionary scenarios to their counterpart model within learning theories. One additional phenomenon relevant to life across all scales is competition for finite resources. Competition takes place at two timescales: on the ecological timescale, species abundances change according to their current ability to access the resources; on the evolutionary timescale, species themselves (i.e., their phenotypic average) become more adapted to the distribution of environmental resources in niche space. Ecological competition therefore serves as an important and, as we will see, the simplest nontrivial example in which combining dynamics at both timescales, referred to as selection and adaptation, is necessary. We model evolutionary-ecological dynamics as follows. Multiple species compete to access environmental resources; the amount of resources individuals of a species extract from the environment determines the abundance of the species. Mean phenotypic trait values of species and their abundances co-evolve. What makes the unfolding of these two dynamics non-trivial is that they are coupled: species adapt to the resources available to them, however, available resources are determined by the trait values of other species as well as their abundances. Abundances, set by the amount of resources species access, change according to the evolution of trait values of all species.

Mathematically, we model the environment as a set of data points *x*_1_, *…, x*_*m*_, each of them corresponding to one unit of resource in resource space at their corresponding location within the niche space. Species *i*’s ability to access resource *x*_*j*_, depending on its trait value *θ*_*i*_, is denoted by *u*(*θ*_*i*_, *x*_*j*_). We refer to *u* as utilization. It is these trait values *θ* that species evolve, and therefore the adaptive trade-off any species face is imposed on their utilization function, ∑_*x*_ *u*(*θ*_*i*_, *x*) = 1, instead of on their fitness. Utilization determines how resources are divided: the share each individual of any species with mean phenotype *θ* receives out of resource *j* is proportional to its utilization at *x*_*j*_, *u*(*θ, x*_*j*_). Consequently, species *i*’s share of resource *j, γ*_*ij*_, is proportional to the product of its utilization *u*(*θ*_*i*_, *x*_*j*_) and its relative abundance *π*_*i*_

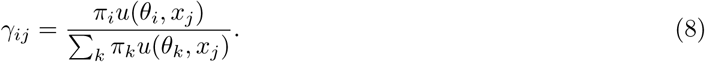

The relative abundance *π*_*i*_ of species *i* is, in turn, given by the total amount of resources its members access,

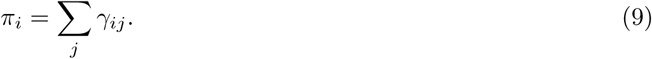

Relative abundances *π* and trait values *θ* co-evolve: relative abundances are updated given more adaptive trait values, whereas trait values evolve given the updated relative abundances. It is worthwhile to investigate how this model relates to a well-known model of ecological competition: the game-dynamical replicator equation. The continuous time game-dynamical replicator equation, describing ecological competition through abundance-dependent selection reads as 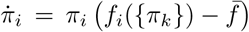, where *π*_*i*_ and *f*_*i*_ is the relative abundance and fitness of type *i*, respectively, and 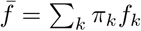 is the average fitness. Importantly, in our model, quasi-equilibrium dynamics is assumed: trait values evolve on a timescale slow enough so that abundances are always in equilibrium given the current trait values. The equilibrium abundance distribution in the game-dynamical replicator equation (if exists), corresponds to all fitnesses being equal to the average fitness, 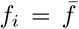 for all types *i*. This can be satisfied by interpreting fitnesses as excess utilization values summed over environmental resources (see Materials and Methods):

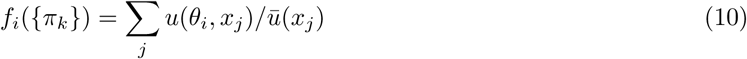

where *ū*(*x*_*j*_) = ∑_*k*_ *π*_*k*_*u*(*θ*_*k*_, *x*_*j*_).

Interestingly, an equivalent formulation of this simple model of evolutionary-ecological dynamics, plays a central role in machine learning and Bayesian statistics: maximum likelihood estimation in mixture models via the expectation-maximization (EM) algorithm. Mixture models fit a weighted sum of distributions *P* (*x*) = ∑_*i*_ *P* (*x*|*{θ*_*i*_*}, h*_*i*_)*π*_*i*_ to the data such that both the parameters of each component distribution *{θ*_*i*_*}* and their relative weights *{π*_*i*_ *}* are optimized in parallel improve on the likelihood of the whole model generating the actual data *x*. Since there is no known general closed-form formula for finding the global maximum of the likelihood function, in practice, mixture models are usually fitted iteratively using a variant of the expectation-maximization algorithm: parameters *θ*_*i*_ are improved given fixed weights *π*_*i*_, whereas weights *π*_*i*_ are improved given fixed parameters *θ*_*i*_; these two steps alternate until the parameters converge (assuming the data *x* does not change). Importantly, both the model framework (mixture models) and the dynamics (EM) are equivalent to the evolutionary-ecological dynamical model described above, with the following translation between the two interpretations: 1. species evolve their trait values *θ*_*i*_ to improve on their utilization function *u*(*θ*_*i*_, *x*) and therefore extract more resources, corresponding to component distributions *P* (*x*|*{θ*_*i*_*}, h*_*i*_) improving on their parameter *θ*_*i*_ to increase the overall likelihood of the model; 2. relative abundances *π*_*i*_ are updated given the amount of resources that is accessible to a species, corresponding to updating the relative weights of the component distribution such that it increases the likelihood of the whole model. Figure 2 illustrates the process.

**Figure 2:**
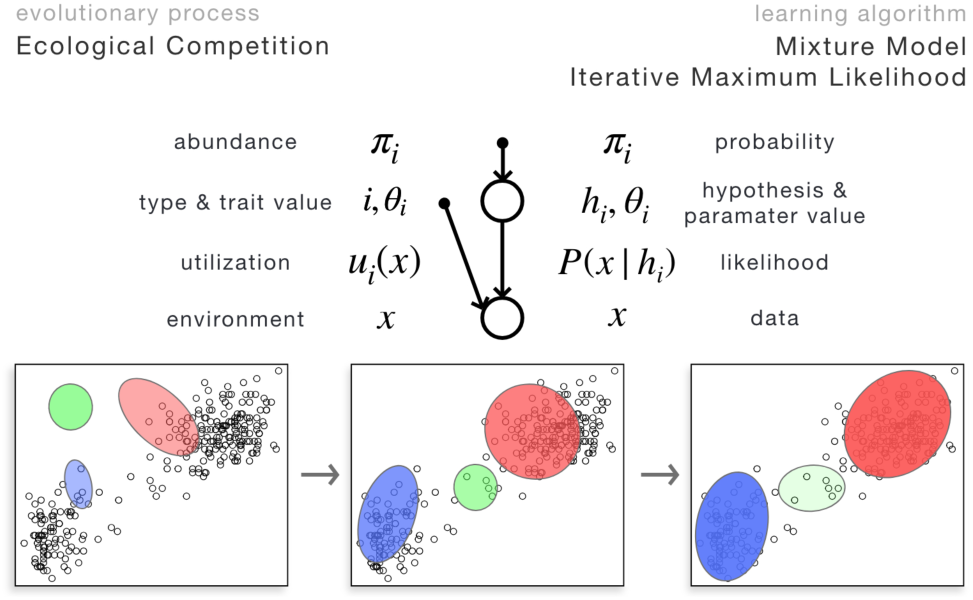
Top: A simple model of evolutionary-ecological dynamics, including intra-species adaptation and inter-species competition for finite resources. Equivalently, this model also corresponds to fitting data with multiple component distributions through optimizing a global likelihood function. This model exemplifies evolutionary systems possessing a global cost (or energy) function that is interpretable in the language of learning algorithms. Bottom: Illustration of evolutionary-ecological dynamics of three species, or equivalently, iterative maximum likelihood (e.g., Expectation-Maximization) optimization of a model involving three component distributions. Each datapoint here corresponds to one unit of resources that is shared between species based on their utilization ability. This resource competition drives species to occupy separate niches, or equivalently, component distributions to avoid overlap. Species abundances (opacity) and their mean phenotypes (location and orientation of the Gaussian contours) co-evolve, driven by the amount of resources they access; equivalently, parameters of component distributions and their weights are iteratively optimized such that the global likelihood of the model is improved.

One important message of this equivalence is that even though it is composed of competing agents, evolutionary-ecological dynamics, in this simple model, optimizes a global cost function, given by the likelihood of the counterpart probabilistic model. Finding a Lyapunov function (i.e., a scalar-valued function over the space of parameters on which the learning/evolutionary dynamics always follows a downwards path) is not always easy, finding one out of many possible ones that contributes meaningfully to a learning-theory based interpretation of emergent computations is even harder. In evolutionary theory, this line of thought goes back to Fisher’s fundamental theorem of natural selection, stating that a scalar function over (phenotypic) parameters, namely, average fitness, always increases if selection is not frequency-dependent [26]. It is perhaps less well known that a Lyapunov function always exists in frequency-dependent scenarios as well, now interpretable in the language of information theory: it is the KL divergence between the current relative abundance distribution over types and the steady state that always decreases over time [27, 11, 28]. Following a similar path to find learning theoretically relevant Lyapunov/potential/cost functions in case of more complex dynamics of competing structures that are capable of representing patterns in high-dimensional data is a potentially important step towards a more unified theory of adaptation, incorporating both evolutionary systems composed of competing representations of the environment and learning systems that learn to represent data, in many cases distributed over myriads of elementary computing units interacting locally. A further fundamental conceptual step would be to account for active manipulation of the environment, allowing causal influences between the evolutionary/learning agent and the environment to be mutual. A work connecting niche construction to active inference [29] nicely exemplifies this idea.

Another message the equivalence of this simple model of evolutionary-ecological dynamics and expectation-maximization in mixture models suggest is that it might be possible to engineer meta-level rules in synthetic evolutionary systems such that they learn as a whole to represent complex patterns in high-dimensional data. A further step towards designing creative artificial systems would be to build such an evolutionary system on complex-enough building blocks so that the complexification of representation can follow millions of diverging paths, impossible to list or foresee all of them by any existing technique, therefore rendering them unexpected and possibly creative.

## Discussion

Bayesian models provide the recipe of performing i) consistent probabilistic computations such that ii) the system extracts the maximum amount of information from the environment given clearly specified model constraints. They therefore form a unique cornerstone of any adaptive dynamics in uncertain environments. Their power lies in their generality: the algorithmic and implementational details constrained by the given physical/informational substrate is not specified. Here we show that it is possible to implement many of the most relevant types of Bayesian computations by simple replicator-based systems. On top of the basic ingredients that include Bayesian update over one hidden variable and iterative optimization of cost surfaces (e.g., maximum likelihood or maximum a posteriori estimation), we describe a replicator-based implementation of many fundamental building blocks of the full Bayesian apparatus, such as inference in mixture models, hidden Markov models and hierarchical models. What makes such a translation table especially appealing is the simplicity of such implementations on the evolutionary side: they do not require introducing fundamental conceptual novelties, instead they correspond to well-known dynamics such as models of ecological competition, mutation and multilevel selection. This observation hints at a more unified picture of evolutionary dynamics and probabilistic learning. It might also provide a ground for transferring more advanced computational techniques from one field to the other. These, we believe, might include leveraging the parallel accumulation of rare subsolutions by recombination, the possibilities of maintained search for novelties by an open-ended replicator system, and, on the side of probabilistic computations, finding an emergent global cost function that is improved upon by the dynamics of competing replicating agents, explaining the seemingly paradoxical adaptive potential of evolutionary systems as wholes.

Another major point is the connection between casting evolution in a stochastic environment as an optimization procedure and Bayesian model selection. Although we mostly discuss within-model adaptation in this paper, we believe that exploring modes of between-model adaptation and evolution is a conceptually rich direction. In particular, we wish to draw attention to two points here. First, Bayesian model selection favors models according to their marginal likelihood, which translates to average fitness in the evolutionary setting. Therefore, whenever marginal likelihood is optimized, average fitness is optimized, and vice versa. This is particularly interesting when models with different structures are compared. One mathematically well-founded evolutionary interpretation we discuss in this paper and elaborate on in detail in [16] is based on the identification of hierarchical models and multilevel selection, giving rise to average fitness-driven transitions in individuality, or, equivalently, marginal likelihood-driven Bayesian model selection. Second, the theory we delineate here does not constrain the form of Bayesian hypotheses/replicator types: they can be models with an intricate structure themselves. In that case, the likelihood/fitness of a hypotheses is the marginal likelihood over its internal structure. This is an additional modelling step we did not take in this paper; we believe, however, that discussing evolvability in stochastic environments could greatly benefit from drawing ideas from bias-variance-like tradeoffs in statistical learning theory.

We envisage three aspects of human inquiry that potentially benefit from this correspondence between evolutionary and probabilistic computations.

First, explaining the adaptive potential of any Darwinian dynamics in Nature: those implemented on a genetic basis, and also those that have not been fully acknowledged, and where the usefulness of replicator-based modeling is questionable at this point, such as memetics [30, 31], Darwinian neurodynamics [32, 33] or quantum Darwinism [34]. Such explanations would mostly be based on either i) finding global cost functions that evolutionary systems emergently optimize or ii) relating the computations performed by the system to probabilistic computations that optimally extract information from external data.

Second, many aspects of human and non-human animal cognition is efficiently modeled by the framework of Bayesian inference and generative models. This is in line with selective pressures favoring optimal extraction of information about the environment that in turn enables action selection that leads to maximum survival and reproduction probability. The possible implementations of Bayesian computations on neural substrates, however, is not fully explored. Here we hint at one possible implementation, based on copying and evaluation of any neural information. We do not posit that all Bayesian computations are implemented this way, instead, we point at the possibility that the effective information extraction provided by Bayesian computations might be efficiently combined with the open-ended novelty generation evolutionary systems are capable of.

This leads to our third point: the possibility of leveraging the combination of evolutionary and Bayesian computations in designing future AI systems to generate creative yet optimally informed solutions/actions in any probabilistic environment. This direction would, in the very first place, need a more thorough understanding of the relation between evolutionary dynamics and learning theories, possibly unified in the same mathematical language. Indeed, besides highlighting parallels between evolutionary dynamics and learning processes, we believe that one of the most important contributions of this paper is the explicit introduction of Bayesian *graphical* models to evolutionary modeling. Bayesian graphical models provide a consistent modular syntax for combinatorial model building, it unloads the cognitive costs of consistency checks and eases building mental maps of related models. We do not state that all evolutionary processes fit into this framework, neither that the Bayesian graphical framework will be the ultimate language of evolutionary modeling, but we strongly believe that a similar combinatorial syntax could aid focusing on relevant aspects of modeling in evolutionary theory.

## Supplementary Information

### The rate of adaptation *η* equals to the phenotypic population variance in the direction of fitness gradient, var(*θ*_||_)

In the following, cov, var, corr and std denote covariance, variance, correlation and standard deviation, respectively. We assume for simplicity that the variance in trait values is small enough to approximate the fitness linearly around the population average by 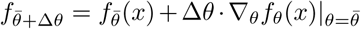, and we also assume for simplicity that the mutant cloud in the phenotype space is isotropic around the population average. Furthermore, it is also assumed throughout the paper that the mapping *f*_*θ*_(*x*) from trait values *θ* and environment *x* to fitness *f* is deterministic, implying that corr(*f, θ*) = 1. Since selection acts in the direction of the (log) fitness gradient ∇_*θ*_ ln *f*_*θ*_(*x*), we project the trait vector *θ* into that direction, denoted by *θ*_||_. Applying the one dimensional Price equation to *θ*_||_ and using the identity |∇_*θ*_*f*_*θ*_(*x*)|std(*θ*_||_) = std(*f*) and 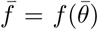 validated by the linear approximation of the fitness landscape, we obtain 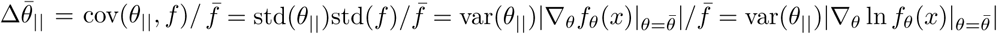.

### The equilibrium condition of the game-dynamical replicator equation can be satisfied by defining fitness as excess utilization

Mathematically, *f*_*i*_(*{π*_*k*_*}*) = ∑_*j*_ *u*(*θ*_*i*_, *x*_*j*_)*/ū*(*x*_*j*_) satisfies the condition of 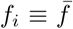. Indeed, by definition, 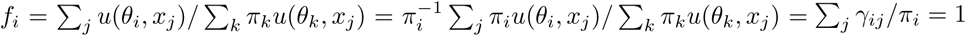 for all *i*.

## Author contributions

D.C., H.G., E.S. designed research, D.C., H.G. performed research, D.C., H.G., I.Z., J.T., E.S. wrote the paper.

## Acknowledgements

We thank Tollevin Williams for his help in producing the figures. The authors acknowledge financial support from the National Research, Development, and Innovation Office under NKFI-K119347 (E.S.), NKFIK124438 (I.Z.), ‘Theory and solutions in the light of evolution’ GINOP-2.3.2-15-2016-00057 (D.C., I.Z. and E.S.), and ‘Learning and evolution’ KKP_19_129848 (D.C. and E.S.); the Volkswagen Stiftung initiative ‘Leben? – Ein neuer Blick der Naturwissenschaften auf die grundlegenden Prinzipien des Lebens’ under project ‘A unified model of recombination in life’ (E.S.); and the Templeton World Charity Foundation under grant number TWCF0268 (D.C. and E.S.).

